# Oxic microbial electrosynthesis can be more energy efficient for biomass production than knallgas or photosynthesis based processes

**DOI:** 10.1101/2023.05.24.542149

**Authors:** Leonie Rominger, Max Hackbarth, Tobias Jung, Marvin Scherzinger, Harald Horn, Martin Kaltschmitt, Johannes Gescher

**Affiliations:** Institute of Technical Microbiology, Hamburg University of Technology (TUHH), Kasernenstraße 12 (F), 21073 Hamburg, Germany; Engler-Bunte-Institut, Water Chemistry and Water Technology, Karlsruhe Institute of Technology (KIT), Engler-Bunte-Ring 9, 76131 Karlsruhe, Germany; Department of Applied Biology, Institute for Applied Biosciences, Karlsruhe Institute of Technology (KIT), Fritz-Haber-Weg 2, 76131 Karlsruhe, Germany; Institute of Environmental Technology and Energy Economics, Hamburg University of Technology (TUHH), Eissendorfer Str. 40 (N), 21073 Hamburg, Germany

**Author notes:** **Contact information for all authors:** Leonie Rominger; Max Hackbarth Tobias Jung Marvin Scherzinger Harald Horn Martin Kaltschmitt.

**Keywords:** Kyrpidia spormannii, Hydrogen oxidizing bacteria, oxic Microbial Electrosynthesis (OMES), Energy Efficiency, Coulombic Efficiency

## Abstract

Data on the efficiency and development of continuous processes are scarce in the emerging field of oxic microbial electrosynthesis (OMES). Therefore, the recently isolated knallgas bacterium *Kyrpidia spormannii* was observed in a bioelectrochemical flow cell setup to study biomass formation and energy efficiency of cathode dependent growth. The study revealed that a potential of -500 mV vs. the standard hydrogen electrode (SHE) caused differences in the structure of the biofilm developing on the cathode, but had almost no impact on biomass growth behavior compared to -375 mV vs. SHE. No growth was observed at 0 mV vs SHE. Coulombic efficiency (CE) was calculated for the cultivation at -500 mV vs. SHE. The process can be conducted with the same electron efficiency as traditional gas fermentation. The solar energy demand with 67.89 kWh kg^-1^ dry biomass is highly competitive to alternative and already established processes for converting (solar) energy to biomass. Additionally, with suggestions for a biomass harvesting method and subsequent recultivation, proof of principle for a continuously operable process was provided. The results pave the way for a new concept in carbon dioxide-based biotechnology.

**Significance statement:** To mitigate global climate change, it is imperative to transition the human economy to a different resource foundation, moving away from fossil fuels and reducing atmospheric carbon dioxide levels. Biotechnological production based on CO_2_ necessitates a supply of energy and electrons. This study reveals that an oxic process, wherein bacteria are directly cultivated as a biofilm on the cathode surface of a bioelectrochemical system, can exhibit higher energy efficiency than plant-based systems or systems reliant on hydrogen generated through water electrolysis. This technology could be instrumental in establishing carbon dioxide and renewable energy as the foundation for feed, food, and platform chemical production.

## 1. Introduction

The fast global carbon cycle relies on the photosynthetic capture of carbon dioxide through oxygenic photosynthesis. Most of the carbon captured by plants from the atmosphere is temporarily stored as biomass and subsequently released back into the atmosphere through oxic processes. A smaller portion becomes embedded underground (and thus removed from the atmosphere) forming fossil resources within geological timeframes. Unfortunately, extensive utilization of these fossil resources since the advent of industrialization, has resulted in the release of fossil-fuel derived carbon back into the fast global carbon cycle. This leads to increased atmospheric CO_2_ concentrations and the consequent climate change, which presents a significant challenge for human development [1]–[4]. Addressing this problem necessitates a transformation in the global economy. A shift is required in the substrate basis used for technical processes towards biobased organic substrates or carbon dioxide. Technical solutions exist for reducing carbon dioxide with electrolysis-derived hydrogen and converting this carbon into long-chain hydrocarbon molecules [5]–[7]. Additional to such heat-induced chemical processes, biological catalysis using basically the same feedstock is also able to provide long-chain hydrocarbons. Compared to thermo-chemical processes, these biological processes typically operate under milder conditions, require less energy, and can direct organisms toward producing a wide range of precisely defined end products [8]–[11].

Reducing carbon dioxide in either of the fixation cycles that evolved in nature necessitates energy and reducing equivalents [12]. In biotechnological applications, hydrogen is currently the most commonly used energy source and electron donor for carbon dioxide-based processes that do not rely on photosynthetic organisms [13]. Oxygenic photosynthesis is widely employed for the latter, but similar to litho-autotrophic carbon fixation with hydrogen, it faces certain challenges. Hydrogen exhibits low solubility in water, which hinders its application in achieving competitive spacetime yields or necessitates significant energy inputs for mixing and homogeneous distribution of small hydrogen bubbles. Contemporary water electrolysis can be utilized for hydrogen production. However, the energy efficiency of this process currently stands at 70 %, and it typically relies on noble electrodes and expensive membranes to separate the anode and cathode compartments [14], [15]. Photosynthesis, on the other hand, requires substantial space and water. The space is determined by the surface area available for installation and adequate illumination. Moreover, if sunlight is supposed to be used directly and light is not produced using for instance LEDs, productivity of such systems will vary significantly based on the latitude.

A relatively recent development in the field of biological carbon dioxide capture and utilization is Microbial Electrosynthesis (MES) [16]. In MES, microorganisms interact with a solid-state cathode, serving as an energy and electron source. Energy and electron transfer can occur directly or with hydrogen produced on the cathode surface [13], [17]. The latter seems to be the case for the organism *Kyrpidia spormannii* which was used in this study. The most effective interaction mode in MES is forming a microbial biofilm directly on the electrode surface [18]. The process can be highly energy efficient since the required voltage is significantly lower compared to water electrolysis [19]. This is due to the biological functionalization of the electrode surface, the constant depletion of hydrogen, the enzymatic catalysis of hydrogen production through direct enzyme-electrode interaction, and/or the direct enzyme-mediated electron import into the organisms’ respiratory chain [20], [21]. Consequently, there is no need to employ expensive electrode materials. If the organisms utilize hydrogen produced on the cathode surface, the process can be directed towards complete depletion during diffusion through the biofilm, effectively circumventing the issue of hydrogen solubility in water through process control.

Currently, most MES processes are carried out by anaerobic organisms, specifically acetogens and methanogens. These anaerobic processes exhibit high electron efficiencies. However, these organisms have limited end-product spectra, low biomass growth coefficients, and genetic engineering can be quite challenging [22], [23]. Additionally, a membrane is required to separate the anode and cathode, posing challenges in terms of reactor construction.

Recently, aerobic bacteria have been introduced into MES processes, offering several advantages. These organisms catalyze a process accompanied by a high energy gain. The increased energy availability provides greater flexibility for genetic engineering of the metabolism, enabling the production of a diverse array of end products. Additionally, biomass growth rates are higher when compared to acetogens and methanogens, and the production of single-cell protein as a viable product appears feasible [24], [25].

A lack of fundamental information regarding the energy efficiency of oxic Microbial Electrosynthesis (OMES) processes hinders their future development. Additionally, if biomass is intended to be a product of an OMES biorefinery, it remains unclear how the biomass cultivated on the cathode can be harvested continuously. To address these questions, this study utilized the recently isolated thermoacidophilic knallgas bacterium *Kyrpidia spormannii* [26], [27]. Biomass growth rates on cathode surfaces were assessed using optical coherence tomography. Coulombic efficiency (CE) was determined and correlated with electrode coverage. The energy efficiency of OMES was compared to hydrogen production based on water electrolysis, followed by biotechnological conversion as well as oxygenic photosynthesis. The results indicate competitive energy efficiency compared to photosynthesis and the two-step process, providing a basis for evidence-based decisionmaking in the development of OMES for future applications in energy conversion, CO_2_ utilization, and capture. Furthermore, this study demonstrates the feasibility of continuous biofilm-based processes on cathodes by employing intermittent shearing of biomass from the electrode by forming hydrogen bubbles.

## 2. Material and Methods

### 2.1 Cultivation

Heterotrophic *K. spormannii* pre-cultures were cultivated in modified R2A-Medium (5 g L^-1^ yeast extract, 1 g L^-1^ peptone, 1 g L^-1^ sodium pyruvate, 0.5 g L^-1^ casein hydrolysate, 0.5 g L^-1^ soluble starch, 0.1 g L^-1^ K_2_HPO_4_, 10 mM MOPS-buffer; pH set to 6.0 with 1 M H_2_SO_4_) at a temperature of 60 °C. ES minimal medium (0.53 g L^-1^ NH_4_Cl, 0.15 g L^-1^ NaCl, 0.04 g L^-1^ KH_2_PO_4_, 0.12 mL L^-1^ MgSO_4_ (1 M), 1 mL L^-1^ CaCl_2_ (0.1 M), 1 mL L^-1^ Wolfe’s mineral elixir; pH 4.0) was used for all electroautotrophic cultivations. The heterotrophically grown pre-cultures were washed in ES medium twice before they were used for inoculating the bioelectrochemical system (BES). A recently described flow cell system was used for bioelectrochemical experiments [28]. For carbon supply, a CO_2_-enriched medium was pumped from a stainless-steel vessel to the biofilms in the flow cell with a flow rate of 100 mL min^-1^. The vessel headspace of 150 mL was pressurized to 1.5 bar overpressure. The system was operated in batch mode. The gas phase was changed every two to three days or according to oxygen demand. The medium was continuously stirred and heated to 60 °C.

The cathode served as the sole energy and electron (hydrogen) donor for the organisms. The electrical energy input was regulated using chronoamperometry. A Gamry Interface 1010 potentiostat (Gamry Instruments, Warminster, USA) was employed for potential control. The potential was applied relative to an Ag/AgCl reference electrode (SE23I, Sensortechnik Meinsberg, Waldheim, Germany). This manuscript references the potential of the standard hydrogen electrode (SHE; 0 mV against SHE corresponds to +199 mV against Ag/AgCl).

### 2.2 Harvesting the biofilm

The biofilms were detached from the electrode surface by applying a two-minute potential pulse ranging from -0.8 V to -2.8 V vs. SHE. This process induced the formation of macroscopic hydrogen bubbles between the cathode surface and the biofilm, leading to the detachment of the biofilm. The detached biomass was subsequently collected from the medium (see details below). If cultivation was to be continued, the entire medium was replaced with fresh, abiotic ES medium before resuming chronoamperometry.

### 2.3 Optical coherence tomography (OCT) and image processing

The development of the biofilm was monitored using Optical Coherence Tomography (OCT) with a Ganymede device equipped with an LSM03 lens (Thorlabs, Dachau, Germany). To ensure consistency and minimize the impact of flow effects on the imaging process, all images were taken at a fixed position in the center of the cathode [28].

A total area of 6 mm × 4 mm was scanned to obtain three-dimensional images. In order to facilitate comparison with other cultivation systems, the normalized biovolume was defined as the volume per unit area (μm^3^μm^-2^) [29]. By deriving the biovolume over time, the accumulation rate (mm^3^ d^-1^ or μm^3^ μm^-2^ d^-1^) could be determined. Processing of OCT images was conducted with ImageJ [28], [30]. In addition to biovolume and accumulation rate, the substratum coverage and roughness coefficient were utilized as additional biofilm characteristics [29]. Biofilm porosity was calculated by subtracting the normalized biovolume from the mean height and then dividing the result by the mean height.

### 2.4 Quantifying the density of the dry biofilm

For energy efficiency calculations, the biovolume was converted into biomass. This conversion involved determining the density of the dry biofilm (ρ_DBM_). To achieve this, the detached portions of the biofilm obtained after harvesting were collected by passing the medium through a sterile filter. Subsequently, the filter was dried for at least 4 days at 70°C. By measuring the difference in weight, the weight of the harvested dry biomass (DBM) could be determined. Establishing a correlation between the difference in biovolume before and after the harvest from triplicate experiments enabled the establishment of a conversion factor of 30.4 ± 2.1 μg mm-^3^.

### 2.5 Determination of H_2_/CO_2_ ratio for autotrophic growth of K. spormannii

In addition to carbon fixation, microbial electron consumption also involves catabolic processes. To determine the actual electron demand of *K. spormannii*, the H_2_/CO_2_ ratio was measured during autotrophic growth. The experiments were conducted using a gas-tight batch system. Initially, a defined gas composition of 70 % H_2_, 20 % CO_2_, and 10 % O_2_ was set using three gas mass flow controllers (Bronkhorst, NL). The gas composition was measured at the beginning and end of the cultivation using a MikroGC 490 (Agilent Technologies). The consumption of H_2_ and CO_2_ was determined based on the difference in concentration. A ratio of 4.23 mol H_2_ (mol CO_2_) was calculated. The corresponding electron efficiency for CO_2_ fixation, which is 53.2%, falls within the range of values determined for knallgas bacteria (*C. necator* 46.21% [31], *H. eutropha* 50.22% [32], *M. fumariolicum SolV* 34.15% [33]).

### 2.6 Calculation of Coulombic Efficiency (CE) and energy demand

Coulombic efficiency (CE) was calculated according to (t in days):

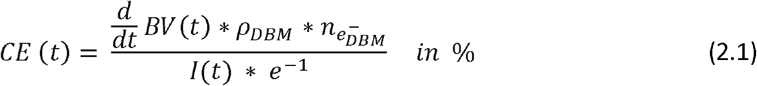

The numerator represents the number of electrons required to form the observed biovolume at any given time during cultivation. BV(t) represents the absolute biovolume development (mm^3^) over time. The change in biovolume during infinitesimally small time intervals around each time point was calculated by taking the derivative concerning time. Using the density ρDBM, the biovolume can be converted into dry biomass, and the number of electrons needed to form 1 mg of dry biomass was determined. In this calculation, biomass was assumed to have a composition of CH_1.8_N_0.18_O_0.38_, corresponding to a carbon oxidation state of -0.5 [34]. In CO_2_, carbon has an oxidation state of +4. Thus, in theory, 4.5 mol of electrons are required to fix 1 mol of CO_2_. Based on the H_2_/CO_2_ consumption ratio of the organism during autotrophic growth with hydrogen as the electron donor and energy source, the actual electron demand (ne-, DBM) was calculated to be 8.46 mol of electrons per mol of CO_2_. The denominator involves calculating the total number of supplied electrons for infinitesimally small time points. This is achieved by dividing the supplied electric current I(t) by the elementary charge e.

To prevent the inclusion of cells from the pre-culture that may falsely appear as autotrophically grown in the system, the medium was replaced with fresh anaerobic ES medium after one day, once the biofilm initiation occurred. This approach ensured that each subsequent increase in biovolume originated solely from cells grown within the system and not from the initial inoculum.

In addition to Coulombic efficiency, the energy efficiency of a process can also be characterized by the biomass yield per supplied energy. The supplied electrical energy was calculated by integrating the electrical capacity input over time. The electrical power was determined by multiplying the electrical current by the average cell voltage (U_cell_).

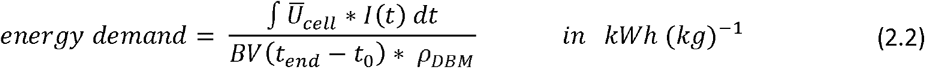

### 2.7 Modeling the oxygen demand with COMSOL Multiphysics

To determine the optimal oxygen concentration in the gas phase, a one-dimensional biofilm model was developed to simulate the growth conditions of *K. spormannii* in an autotrophic OMES. The model was constructed using COMSOL Multiphysics® 6.0 (2021). The principle is based on a model developed by Korth et al. [35], but instead of anodic biofilm growth and direct electron transfer, a cathodic biofilm based on hydrogen as an electron shuttle was modeled. Also, the model used in this manuscript is less complex. It depicts the concentration profiles of oxygen and hydrogen across the thickness of the biofilm at a specific location. The concentration of CO_2_ was not considered since its solubility is relatively high, suggesting it is unlikely to be limiting in the developed process. Oxygen and hydrogen concentrations within the biofilm were determined by subtracting consumption effects from diffusion. The two substrates diffused through the biofilm in a counterdiffusion manner. The boundary conditions for the model included the dissolved oxygen concentration in the bulk phase and the hydrogen evolution rate at the substratum, which were derived from experimental data (Table 1). The growth rate of the biofilm was calculated using Monod kinetics for both substrates. All relevant formulas and parameters can be found in Table 1. The model was simulated for a cultivation temperature of 60°C.

**Table 1:**
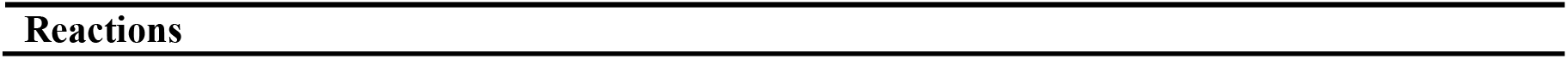
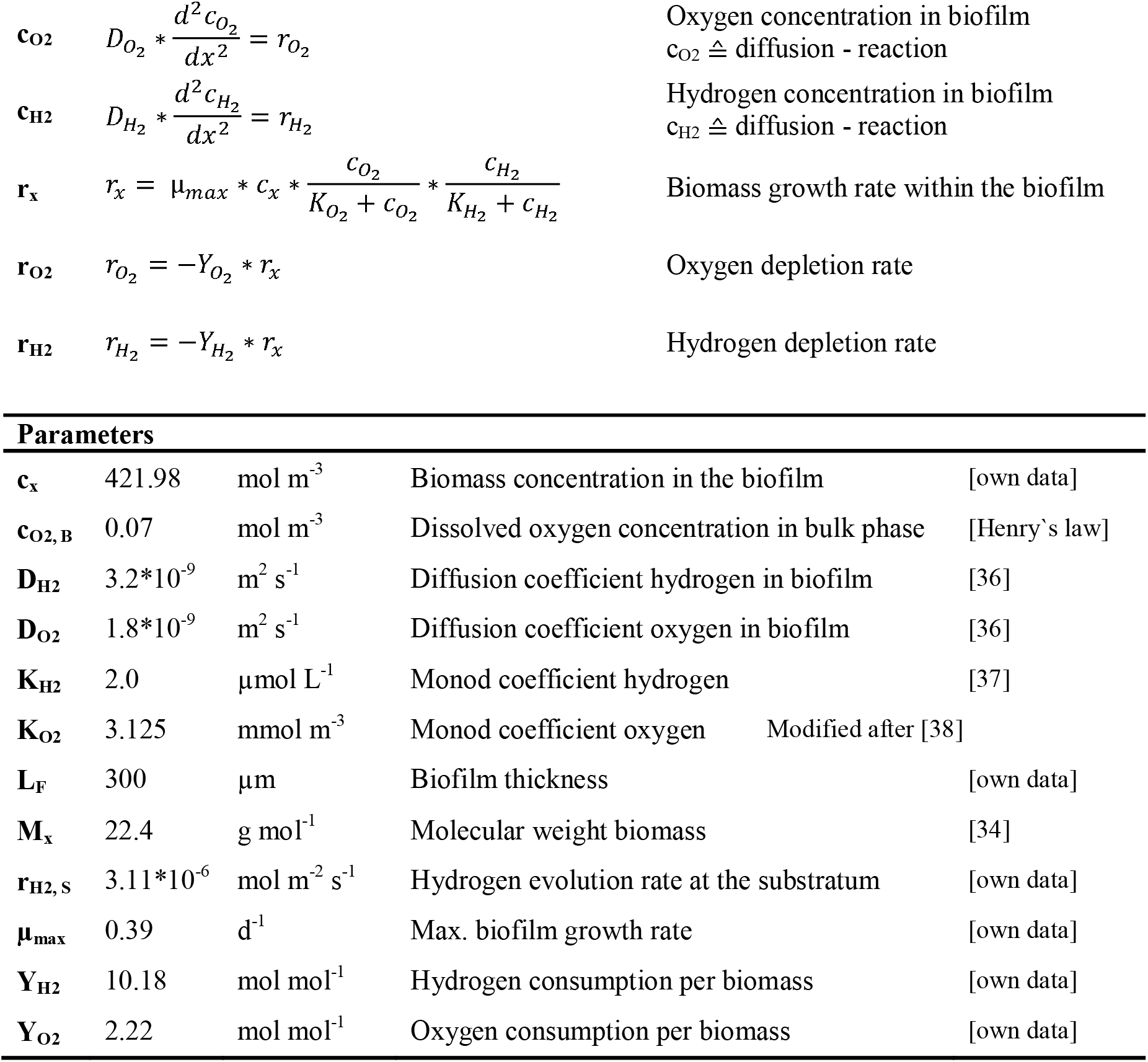
Formulas for calculation of stationary substrate concentrations (oxygen and hydrogen) and biofilm growth rate over the length of the biofilm; all parameters used for the model are listed.

|  |  |  |
| --- | --- | --- |
| **c<sub>O2</sub>** | $D_{O_2} * \frac{d^2 c_{O_2}}{dx^2} = r_{O_2}$ | Oxygen concentration in biofilm<br>$c_{O_2} \triangleq$ diffusion - reaction |
| **c<sub>H2</sub>** | $D_{H_2} * \frac{d^2 c_{H_2}}{dx^2} = r_{H_2}$ | Hydrogen concentration in biofilm<br>$c_{H_2} \triangleq$ diffusion - reaction |
| **r<sub>x</sub>** | $r_x = \mu_{max} * c_x * \frac{c_{O_2}}{K_{O_2} + c_{O_2}} * \frac{c_{H_2}}{K_{H_2} + c_{H_2}}$ | Biomass growth rate within the biofilm |
| **r<sub>O2</sub>** | $r_{O_2} = -Y_{O_2} * r_x$ | Oxygen depletion rate |
| **r<sub>H2</sub>** | $r_{H_2} = -Y_{H_2} * r_x$ | Hydrogen depletion rate |

| **Parameters** |  |  |  |  |
| --- | --- | --- | --- | --- |
| **c<sub>x</sub>** | 421.98 | mol m<sup>-3</sup> | Biomass concentration in the biofilm | [own data] |
| **c<sub>O2, B</sub>** | 0.07 | mol m<sup>-3</sup> | Dissolved oxygen concentration in bulk phase | [Henry`s law] |
| **D<sub>H2</sub>** | 3.2*10<sup>-9</sup> | m<sup>2</sup> s<sup>-1</sup> | Diffusion coefficient hydrogen in biofilm | [36] |
| **D<sub>O2</sub>** | 1.8*10<sup>-9</sup> | m<sup>2</sup> s<sup>-1</sup> | Diffusion coefficient oxygen in biofilm | [36] |
| **K<sub>H2</sub>** | 2.0 | μmol L<sup>-1</sup> | Monod coefficient hydrogen | [37] |
| **K<sub>O2</sub>** | 3.125 | mmol m<sup>-3</sup> | Monod coefficient oxygen | Modified after [38] |
| **L<sub>F</sub>** | 300 | μm | Biofilm thickness | [own data] |
| **M<sub>x</sub>** | 22.4 | g mol<sup>-1</sup> | Molecular weight biomass | [34] |
| **r<sub>H2, S</sub>** | 3.11*10<sup>-6</sup> | mol m<sup>-2</sup> s<sup>-1</sup> | Hydrogen evolution rate at the substratum | [own data] |
| **μ<sub>max</sub>** | 0.39 | d<sup>-1</sup> | Max. biofilm growth rate | [own data] |
| **Y<sub>H2</sub>** | 10.18 | mol mol<sup>-1</sup> | Hydrogen consumption per biomass | [own data] |
| **Y<sub>O2</sub>** | 2.22 | mol mol<sup>-1</sup> | Oxygen consumption per biomass | [own data] |

The optimal oxygen concentration in the bulk phase was reached if after diffusion through the biofilm the oxygen was almost depleted and abiotic reaction with the cathode was omitted. The hydrogen concentration was defined from the hydrogen evolution rate, which derived from the electric current flowing into the system.

## 3. Results

### 3.1 Growth kinetics of K. spormannii biofilms on cathode surfaces at different applied potentials

First, the effect of the applied potential on biofilm growth and formation was investigated. OCT was used to quantify biovolume. Figure 1(a) shows the relative biovolume, where the biovolume at each time point was normalized to the maximum biovolume of the respective cultivation. The absolute maximum biovolume ranged between 125 and 175 μm^3^ μm^-2^ for all cultivations (Figure 1(d)). The data points of each growth curve were fitted with a polynomial (R > 99 %). Cells did not grow at 0 mV vs. SHE; however, they remained viable even after more than two weeks at this potential since a switch to -500 mV led to the rapid growth of the organisms on the cathodes. Exponential growth commenced immediately after inoculation at -375 mV (green) and -500 mV vs. SHE (blue). The maximum biovolume was reached after approximately eight days. Moreover, in terms of accumulation rate (Figure 1(b)), which represents the derivative of biovolume over time, only minor differences were observed. The maximum accumulation rate at -500 mV vs. SHE was observed after 1.8 days, with a value of 62.05 mm^3^ d^-1^, slightly higher than the value of 52.12 mm^3^ d^-1^ observed at - 375 mV vs. SHE after 3.4 days. For all potentials, the maximum accumulation rate was correlated with complete substratum coverage (Figure 1(c)).

**Figure 1:**
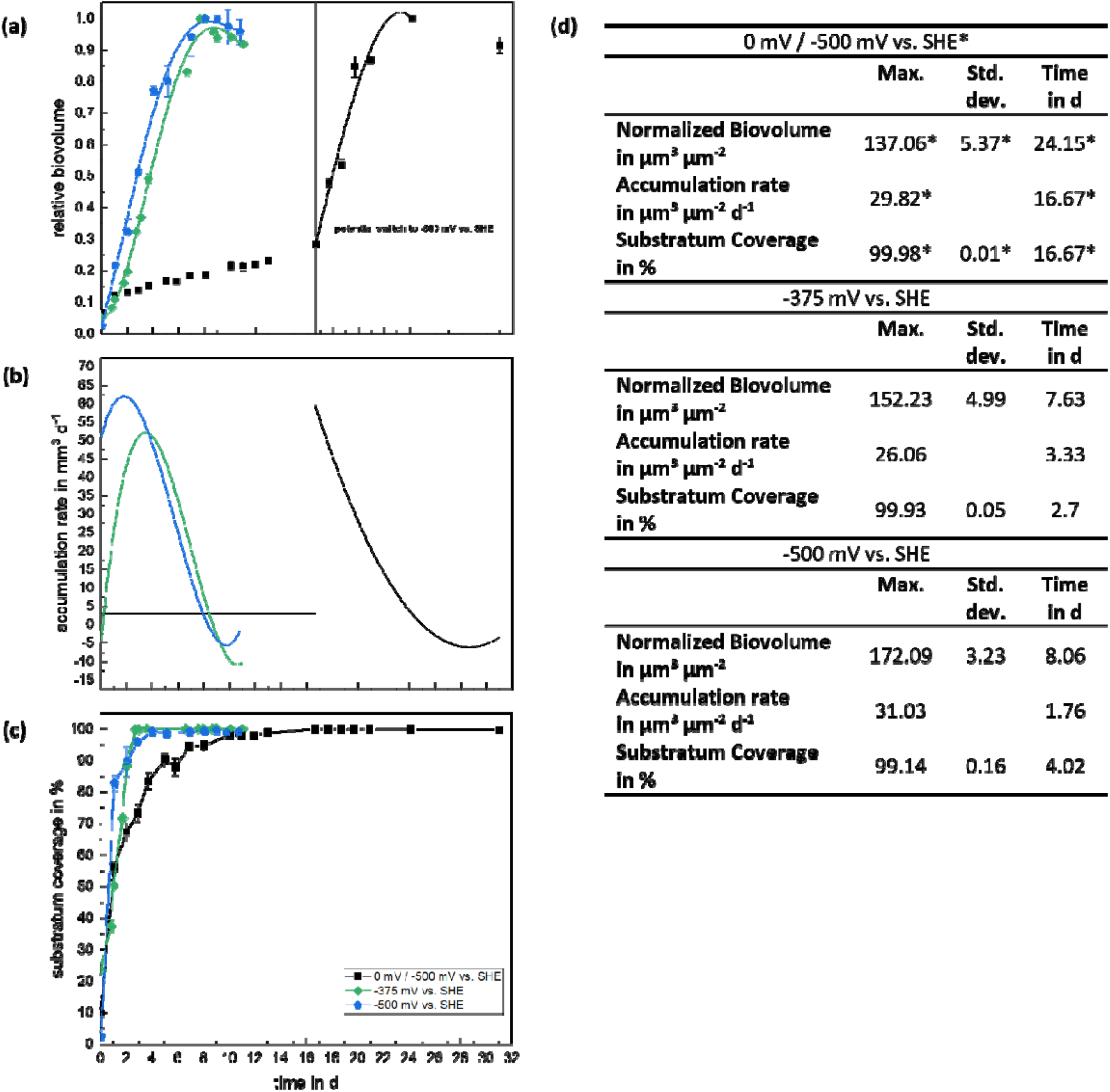
Effects of applied potential on (a) relative biovolume, (b) accumulation rate and (c) substratum coverage; Comparison between growth at -500 mV vs. SHE (blue), -375 mV vs. SHE (green) and 0 mV vs. SHE (which was switched to -500 mV vs. SHE after 16.7 days, respectively) (black). (d) shows the maximum values and standard deviation for normalized biovolume, accumulation rate and substratum coverage, as well as the timepoint; for 0 mV vs. SHE, values that appeared after the switch to -500 mV vs. SHE are indicated with a *.

The biofilm morphology varied among the biofilms grown at different potentials. Figure 2 illustrates the biofilm height maps for mature biofilms. To further characterize the biofilm structure, porosity and roughness were calculated using the same potentials as in the previous studies. At -500 mV vs. SHE, the biofilms exhibited tower-like structures with dimensions of up to 0.3 mm on the substratum surface. The biofilm had a relatively high porosity, with individual tower-like structures separated by an average distance of 0.5 to 1 mm from each other. Similar biofilm structures were observed when the potential was shifted from 0 mV vs. SHE, where no biofilm growth was observed, to -500 mV vs. SHE. This finding supports the notion that this growth pattern is robust at this potential. At -375 mV, representing lower energy availability, the biofilms exhibited denser growth and lower porosity.

**Figure 2:**
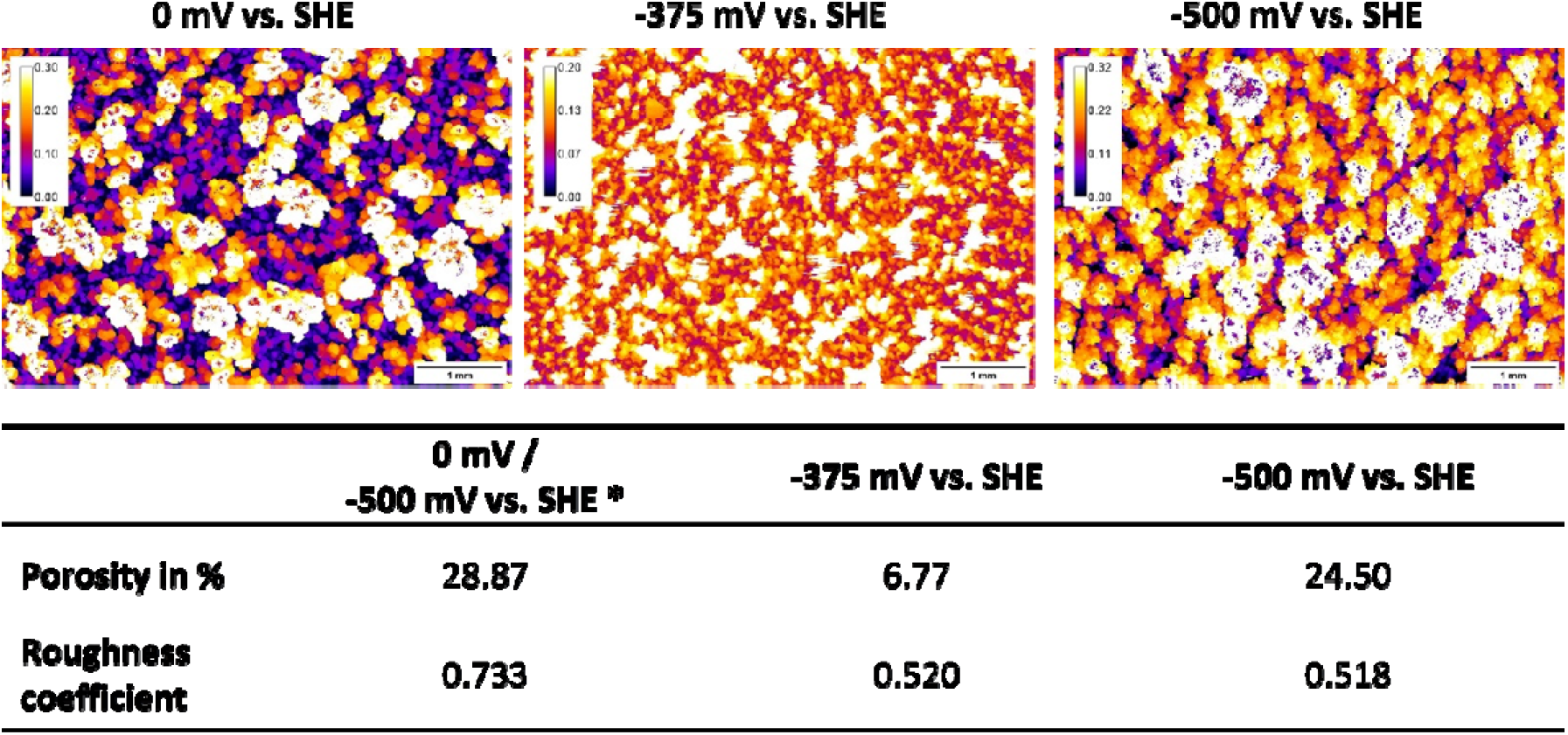
Comparison among biofilms grown at different potentials. Biofilm heightmaps are shown, as well as biofilm porosity and roughness as structure describing parameters. Numbers in the color calibration bar are given in mm. The images were taken on day 31 (0 mV; day 14.33 at -500 mV), day 11 (−375 mV) and day 10.77 (−500 mV), respectively.

### 3.2 Developing a continuous process for biofilm harvest and regrowth

At a certain thickness, biofilm growth is hindered by mass transfer limitations, indicating the point at which biomass should be harvested and used directly as a product or as a substrate in a biorefinery process [38]. The developed method for controlled biofilm detachment is based on the generation of hydrogen at the cathode surface through the application of a highly negative potential. In preliminary abiotic tests, the initiation of visible hydrogen bubble formation was determined by observing abrupt decreases in current density, along with the detection of hydrogen bubbles using OCT imaging (data not shown). Hydrogen bubbles began to form at -550 mV vs. SHE at the chosen process conditions (60 °C, pH 4.0, graphite electrode).

The hydrogen bubbles formed beneath the biofilm, causing partial destruction of the biofilm structure above (Supplementary Figure 2). To quantitatively evaluate the impact of hydrogen bubbles on biofilm detachment, a series of experiments were conducted, gradually decreasing the potential from -0.8 V to -2.8 V. The relative biovolume and substratum coverage were measured using OCT (Figure 3(a)). Between -1.0 and -1.4 V, some residual bubbles cast shadows on the biofilm in the OCT images, leading to an underestimation of the biofilm measurements. Therefore, the presumed more realistic development of biovolume and coverage in this area is depicted with dotted lines in Figure 3.

**Figure 3:**
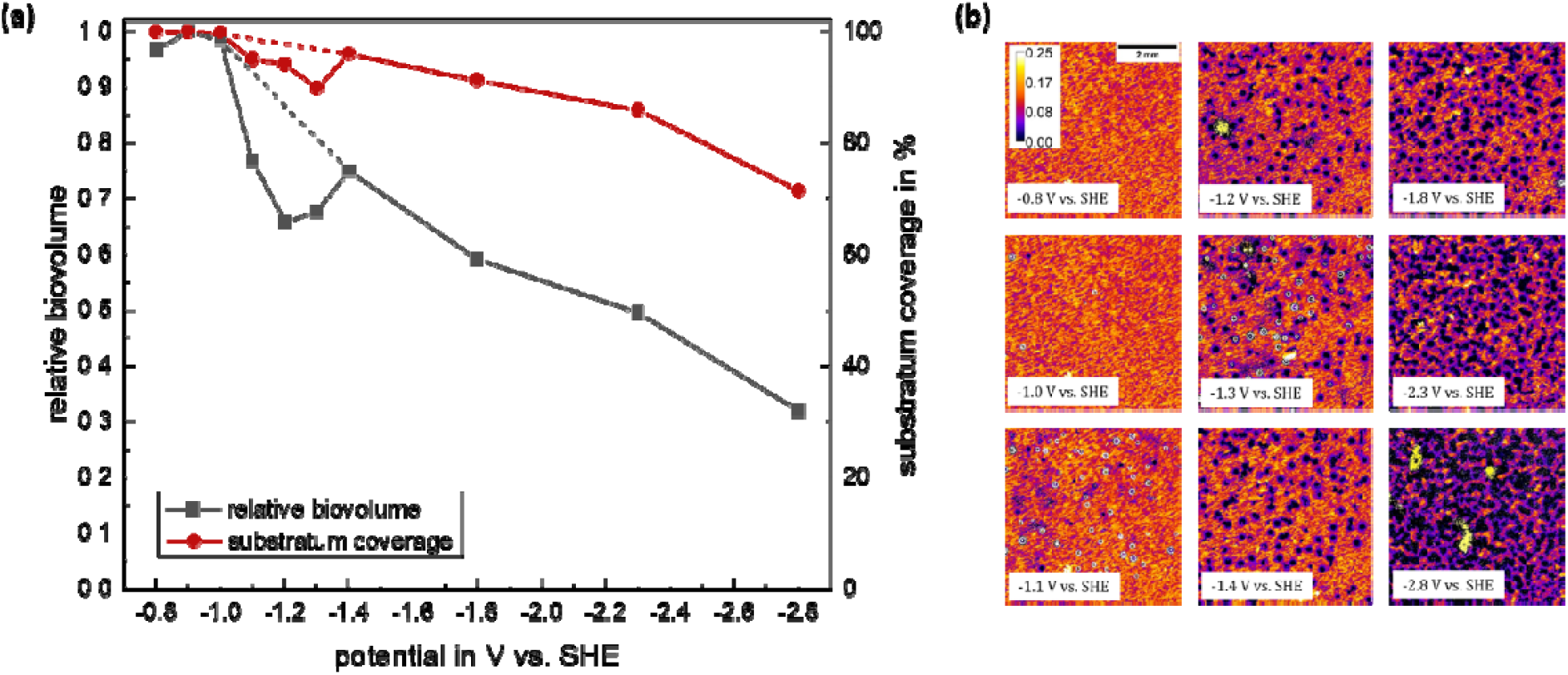
**(a)** Characterization of biofilm detachment by hydrogen bubbles with regards to relative biovolume and substratum coverage. Dotted lines represent the presumable development of biovolume development which was cumbersome to analyze due to hydrogen bubble localization. **(b)** Biofilm height profiles for increasing negative potential; color calibration bar is given in mm

Both the biovolume and substratum coverage exhibited an almost linear decrease with increasing potential. The biovolume decreased by 65%, while the coverage of the cathode surface was ultimately only 70%. Figure 3(b) displays the height profiles of the biofilm during the detachment steps. The heightmaps reveal hydrogen bubble-induced voids within the biofilm, with an approximate radius of 300 μm, starting from -1.1 V vs. SHE. As the potential decreases further, the number and size of these voids progressively increase.

To ensure the feasibility of a continuous process, it is important to demonstrate the biofilm’s ability to regenerate from the remaining cells located on or near the area of bubble formation on the cathode. To investigate this, the medium and all planktonic cells were replaced with fresh medium, and the cultivation was restarted. The resulting biovolume and substratum coverage are presented in Figure 4(a).

**Figure 4:**
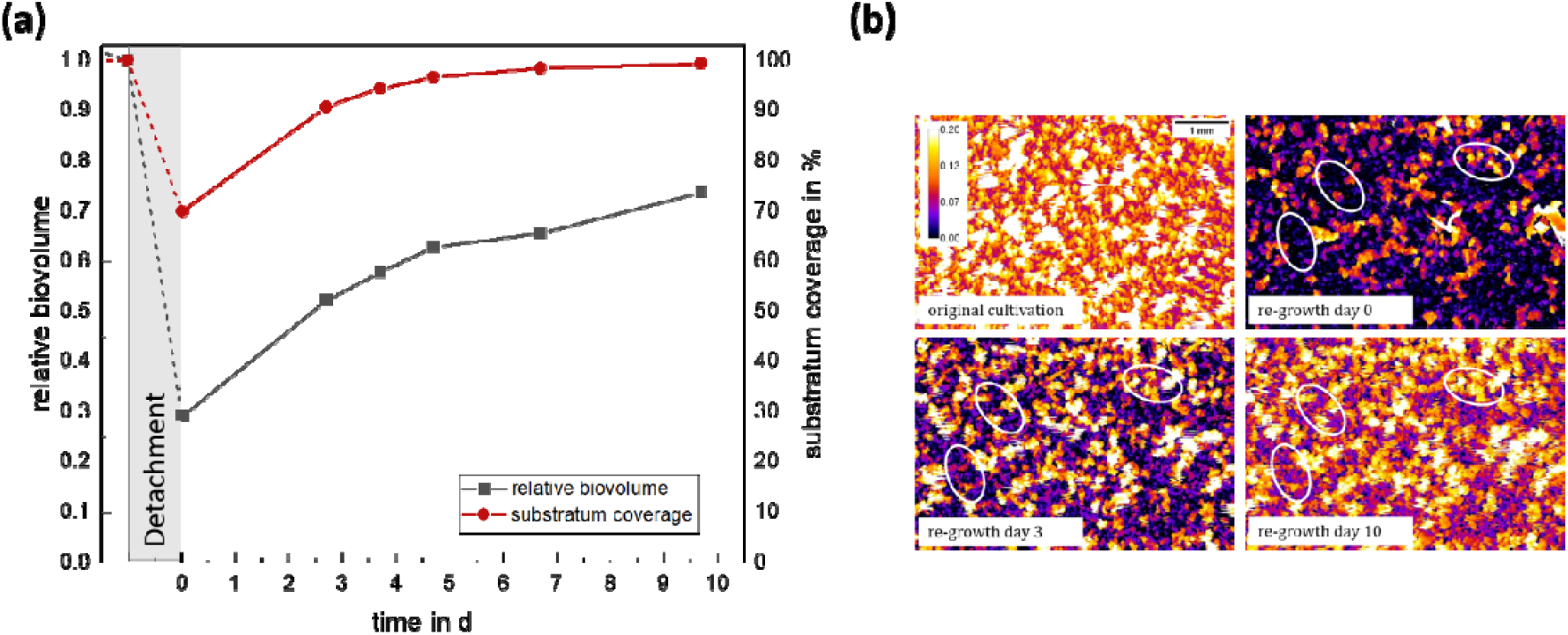
**(a)** Evolution of biovolume and substratum coverage in a harvested system without planktonic cells; **(b)** Heightmaps of the biofilm growing again; color calibration bar is given in mm. Three positions have been marked to allow for better assessment of biofilm regrowth. The two positions on the left are completely uncovered in the beginning. On day ten, the spots can be found covered with a roughly 0.07 mm thick biofilm again. The right marking shows a small biofilm structure with a height of 0.1 mm in the beginning. On day ten, it is 0.15 mm high. By this, the biofilm growth can be followed on uncovered cathode surface, as well as on already existing structures.

After detachment, 30 % of cathode surface were not covered by biofilm anymore. Within the next 10 days, the growing biofilm was able to completely cover the electrode surface again and could regenerate to more than 70 % of its initial volume before detachment (Figure 4 (b)).

### 3.3 H_2_/CO_2_ ratio and electron efficiency for cathode- and knallgas-based growth

The theoretical H_2_/CO_2_ ratio for converting CO_2_ into biomass is 2.25 mol mol^-1^. This is because the carbon in CO_2_ has an oxidation state of +4, which needs to be reduced to -0.5 in biomass. The electron demand for this process is described by the CE of CO_2_ reduction to biomass (Figure 6). However, the actual H_2_/CO_2_ ratio must be higher due to catabolic processes. The thermodynamic lower limit for the H_2_/CO_2_ ratio for autotrophic growth was calculated to be 2.4 mol H_2_/mol CO_2_ [39], [40]. For the organism K. spormannii studied here, the actual H_2_/CO_2_ ratio was determined to be 4.23 mol mol^-1^. Achieving a similar electron efficiency in an OMES process compared to gas fermentation in a batch system is challenging because abiotic oxygen reduction at the working electrode can act as a significant electron sink. To mitigate this risk, we developed a model for the OMES process that considers metabolic data from the litho-autotrophic growth of K. spormannii, as well as cathodic hydrogen production and the diffusion of oxygen and carbon dioxide from the bulk phase into the biofilm. Figure 5 shows the concentration profiles for the adjusted conditions (4 % oxygen in the pressurized gas phase; hydrogen evolution rate of 3.11 10^−6^ mol (s m^2^)^-1^). The graph also illustrates the deduced growth rate, r_x_, in red.

**Figure 5:**
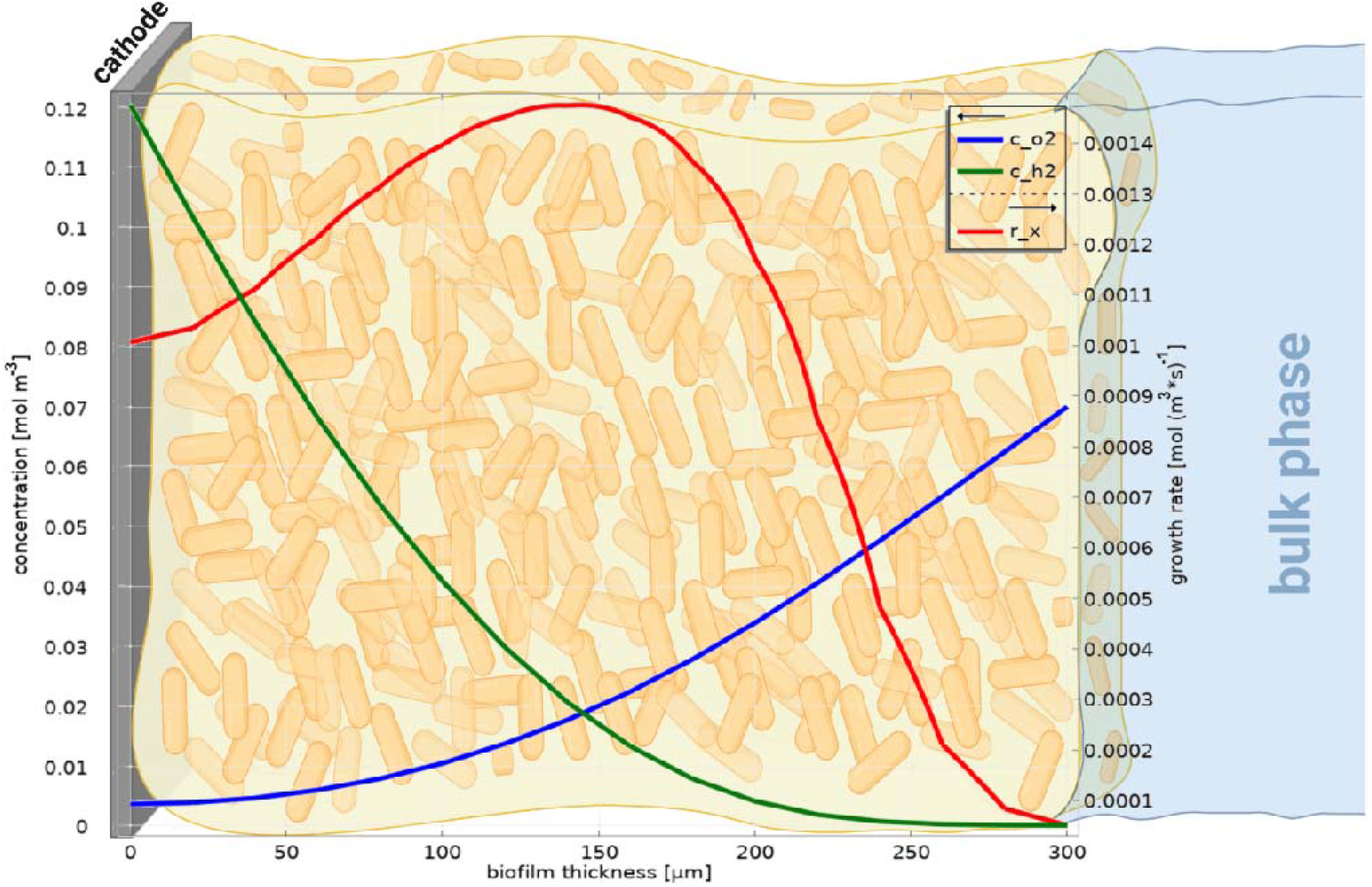
Concentration profiles of oxygen (blue) and hydrogen (green) among the biofilm height; in red, the growth rate is plotted.

**Figure 6:**
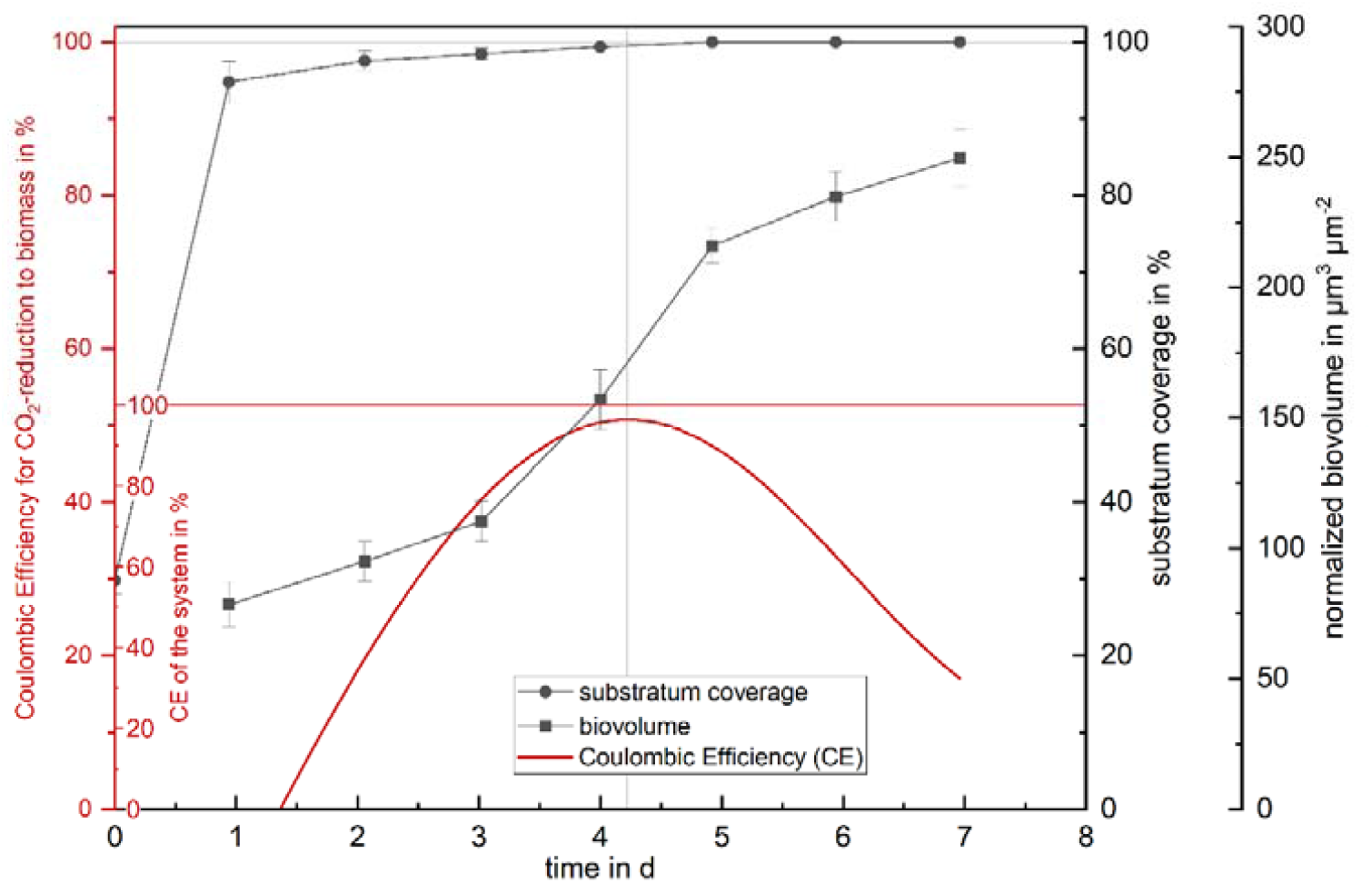
Coulombic Efficiency (CE; red), normalized biovolume (black squares) and substratum coverage (black circles) over time for biofilm growth at ⍰500 mV vs. SHE; CE of the system describes the percentage of electrons that was used for biological processes (anabolism and catabolism), while CE for the reduction of CO_2_ tells how many electrons were used for CO_2_ reduction to biomass (anabolism only).

Using this model, the oxygen concentration was adjusted according to the biovolume during the time course of the experiment, and a working electrode potential of -500 mV vs. SHE was applied. The choice of the relatively low potential of -500 mV was based on the observation that surface coverage occurred most rapidly at this potential, suggesting that abiotic electron consumption would be minimized.

As depicted in Figure 6, the OMES process achieved a biotic electron consumption of up to 95% compared to the knallgas experiment. This demonstrates the successful minimization of abiotic losses (5 %) by adjusting process conditions. The peak CE aligns with the highest rate of biomass accumulation. During this period of maximum CE, the electrode is nearly entirely covered with biofilm. On average, the CE reached 49.51% compared to the knallgas experiment, with the lag phase and stationary growth phase exhibiting lower efficiency compared to the exponential growth phase. The latter is probably due to abiotic oxygen reduction and mass transfer limitation, respectively.

### 3.4 Energy demand of OMES

As one of the primary objectives of this process is to convert carbon dioxide into a valuable product through biomass production, the solar energy required to generate a specific biomass quantity was calculated for the OMES process. A comparison was made to phototrophic biomass production and biomass production based on K. spormannii growing on knallgas using electrolysis-produced hydrogen. For simplicity, it was assumed that only the availability of electrons and energy limited the productivity of the biosystems. The energy demand for the production of 1 kg dry biomass was calculated using the cell voltage, multiplied by the integral of the electric current (s. eq. 2.2). The photovoltaic conversion efficiency of solar energy into electricity was assumed to be 17.5%, a value reported in the literature for monocrystalline silicon solar cells [42]. For the calculation of the OMES process only carbon dioxide-based biofilm growth was considered. Hence, only biofilm growth after medium replacement was evaluated. The median solar energy demand for cathodic growth was calculated to be 210.99 kWh kg^-1^, with a standard deviation of 6.39 kWh kg^-1^. Of note, energy demand calculation for the individual growth stages throughout the experiment displayed in Figure 7 demonstrates the efficiency potential of the technology as the phase with the highest CE corresponds to the lowest solar energy demand of 67.89 kWh kg^-1^. Without considering the photovoltaic conversion efficiency into account, the actual energy demand for this process phase was 11.88 kWh kg^-1^.

**Figure 7:**
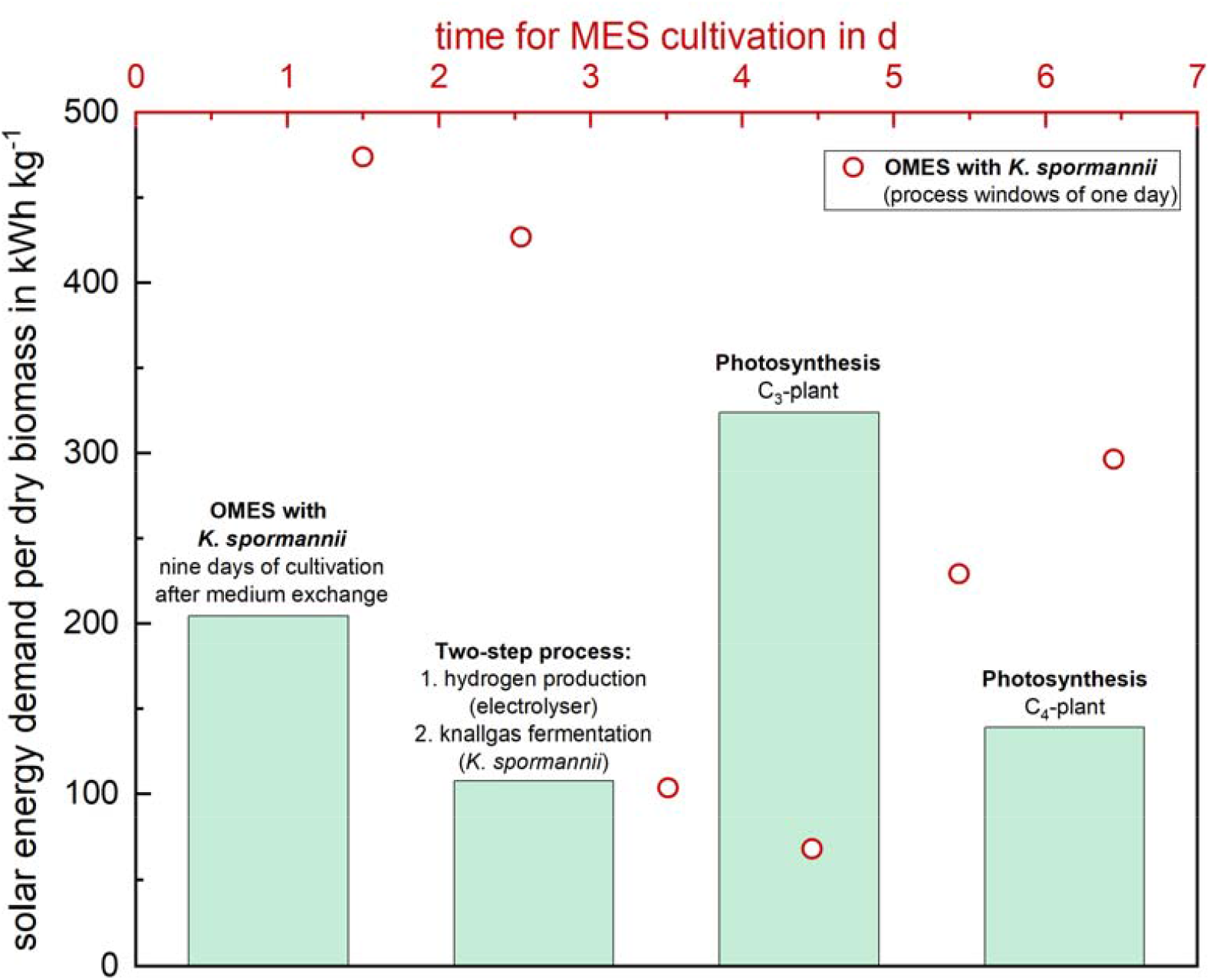
Solar energy demand for the production of 1 kg dry biomass with the cultivation of K. spormannii in the described OMES-process at -500 mV (the whole cultivation period of nine days is considered for the column, single process windows of one day each are considered for the red circles), the production of hydrogen for cultivation of knallgasbacteria (K. spormannii) in an external electrolyser and photosynthesis in plants (C_3_ and C_4_-metabolism)

For knallgas bacteria, which utilize hydrogen as an electron donor, there is also the option to produce hydrogen in an external electrolyzer first, which can then be supplied to the bacteria in a second step. The carbon content in dry biomass accounts for 53.5% of its weight [34]. With a theoretical hydrogen demand of 4.5 mmol H (mmol C)^-1^, a final demand of 0.2 kg of H_2_ kg^-1^ of dry biomass can be assumed. The voltage efficiency for PEM electrolysis and alkaline electrolysis, for cell voltages ranging between 1.8 and 2.24 V, ranges between 62% and 82%. This leads to an energy demand of about 45–50 kWh kg^-1^ of H_2_ [14], [43]. Considering the photovoltaic conversion efficiency of 17.5%, this biomass production method results in a solar energy demand of 57.43 kWh kg^-1^ of dry biomass under the best possible conditions. However, when considering the realistic values for hydrogen efficiency determined in this study, the energy demand nearly doubles to 107.91 kWh kg^-1^ (Figure 7). The OMES and knallgas-based processes were also compared to photosynthesis in C3 and C4 plants. The average conversion efficiencies from sunlight to biomass were estimated to 1.5 % for C3 plants and 3.5 % C4 plants, respectively (these values correspond to the literature data for plants in fields with sufficient nutrient and water supply during their active vegetation phase) [44]. The calorific value for a whole plant was determined to be 0.0175 MJ g^-1^ by Lieth et al. [45], resulting in a solar energy demand of 324.07 kWh kg^-1^ for C3 plants or 138.89 kWh kg^-1^ for C4 plants. Compared to photosynthesis, the process described here can be up to 2.05 times more efficient under the assumptions made (i. e. an exclusive consideration of the energy required in the conversion of radiation to biomass without upstream and downstream process steps). This is primarily due to the higher conversion efficiency of photovoltaic systems compared to photosynthesis [23], [46].

## 4. Discussion

This study aimed to address questions regarding the energy demand for OMES processes using carbon dioxide as a carbon source and identify strategies for biomass harvesting from the cathode in a continuous process. For this purpose, *K. spormannii* was selected as the model organism as previous studies revealed robust growth of the organism on cathodes, allowed bioinformatic analysis of metabolic potential and introduced an experimental infrastructure to measure necessary parameters.

Biofilm growth was not observed at 0 mV vs. SHE, which is not surprising considering that hydrogen is assumed to be the electron donor during cathodic growth. While the exact overpotential for H_2_ production on the applied graphite electrode is unknown, based on process conditions and the Nernst equation, the hydrogen evolution potential can be assumed to be more negative than -102.5 mV vs. SHE. However, the growth limitation was overcome upon switching the potential from 0 to -500 mV, and the biofilm began to grow. This observation confirms that the biofilm can recover and continue growing even after a period of starvation lasting at least 16 days. This characteristic may prove advantageous for future applications, particularly in scenarios where the availability of renewable energy fluctuates.

The applied potential is a key parameter for the performance of an oxic MES. It has an impact on the biofilm structure and influences electrons transfer mechanisms in the biofilm [20], [47], [48]. In this study, similar biofilm growth kinetics were observed at -375 mV and -500 mV vs. SHE, respectively. Applying a potential more negative than -375 mV vs. SHE did not accelerate growth any further. Moreover, the accumulation rate was not affected by the cathode potential. Notably, the time points of maximum accumulation rate were found to correlate with 50 % of the maximum biovolume in the cultivations at -375 mV and -500 mV vs. SHE. Here, hydrogen diffusion from the electrode or oxygen diffusion from the bulk phase may be limiting if a certain biofilm thickness is reached. This biofilm height possibly represents a process limiting parameter, after which biofilm growth is slowed down. A similar observation could be made in a mathematical model of an MES process provided by Li et al. [49]. The difference in biofilm structure related to electrode potential might be due to inhomogeneous hydrogen formation on the cathode surface correlated to lower electrode potentials that will favor the formation of larger hydrogen amounts. To this end, it would be interesting to observe how hydrogen forms at different potentials and what the impact of very small hydrogen bubbles might be on the process. Unfortunately, this was below the resolution limit of the analytic infrastructure and will be a research field in the future.

Mishra et al. calculated an energy demand of 29.24 kWh kg^-1^ biomass for hydrogen oxidizing knallgas bacteria which were fed with externally produced hydrogen [25]. More than 50 % of the supplied electrons were utilized to reduce CO_2_ to biomass. This value is significantly higher than the average anabolic hydrogen utilization of acetogenic or methanogenic organisms, which is approximately 10%. Consequently, biomass production by these organisms carries a significantly higher energy demand due to a very low CO_2_ fixation into biomass conversion efficiency [50]. In fact, according to Mishra et al., the energy demand for autotrophically grown acetate-producing bacteria is 231.48 kWh kg^-1^ biomass [25]. This might also hamper production of other products besides biomass, as biocatalyst regrowth might be a process limitation [23]. Consequently, cathodic biomass production will most likely be more efficient using K. spormannii and comparable organisms. Not only regarding the share of supplied electrons ending up in biomass, but also regarding overall energy efficiency it might be worth considering oxic cathodic processes for future processes, as less energy is necessary for biomass production compared to phototrophic growth, as well as growth based on hydrogen produced in an electrolyser. For the latter, it should also be considered that hydrogen solubility is a main factor limiting space time yields and influencing energy input during a gas fermentation process. In the case of K. spormannii, cathodic growth is biofilm dependent, and hydrogen can be depleted during diffusion through the biofilm matrix. It will be important to design biofilm detachment and regrowth in a way that biofilm development will remain in an optimal process window. The described method of biofilm detachment not only allows for steering process efficiency but also serves as a significant downstream processing step for biomass harvesting. To this end, regarding the current development of global population it seems inevitable to reconsider the way how protein for food and feed is produced. For this so called “protein transition” the here described process might be an option.

The dissolved oxygen concentration is an important factor for optimizing efficiency. In the initial growth phase, there are presumably higher energy losses due to reduction of oxygen to reactive oxygen species. With progressing surface coverage, less cathode surface is available, thus less oxygen can be reduced abiotically. By eliminating the oxygen reduction sink at the cathode, the anodically produced oxygen can likely supply the biofilm. This observation aligns with the higher oxygen demand during the phase until full substratum coverage is achieved, compared to later process stages requiring minimal oxygen feed. Optimizing the oxygen concentration based on the biofilm’s current growth state makes it possible to minimize energy losses.

Considering all these factors, there is significant potential for energetically optimizing the energy efficiency of the OMES process. In addition to the applied cathode potential, the oxygen concentration, biofilm thickness, density, and reactor design will also play critical roles. Selecting the appropriate electrode material and size can contribute to reducing cell voltage and thus achieving the desired energy input [23], [51], [52]. Furthermore, genetic engineering holds promise for further enhancing efficacy. For instance, Li et al. demonstrated that introducing a cyanobacterial RuBisCO into *Cupriavidus necator*, another knallgas bacterium, resulted in improved autotrophic growth and production of polyhydroxybutyrate (PHB) [52]. Similar strategies could potentially advance the performance of *K. spormannii* to a similar extent.

## 5. Conclusion

The primary objective of this study was to overcome the challenges that have thus far hindered the efficiency of MES processes. According to Prévoteau et al. [23], these challenges predominantly include non-selective production at low product titers, low production rates, high cell voltage (ohmic drop) resulting in low energy conversion efficiency, and high capital costs. Our findings demonstrate that the cathode potential between -375 mV and -500 mV vs. SHE does not affect the biofilm growth rate of *K. spormannii* but does influence the biofilm structure. By applying a highly negative potential, we propose a method for partially harvesting the biofilm by generating hydrogen bubbles. The biofilm can regenerate after partial harvesting, enabling continuous operation of the process. The CE for biobased electron usage reaches nearly 100 % at its maximum, surpassing the energy demand of oxygenic photosynthesis and knallgas-based processes for biomass production from CO_2_. Moreover, oxic cathodic processes eliminate the need for a membrane to separate the anode and cathode, resulting in a substantial decrease in capital costs, as membranes typically constitute the most expensive component of bioelectrochemical systems.

## Appendix

### Abbreviations

BV: Biovolume
CE: Coulombic Efficiency
DBM: Dry Biomass
OCT: Optical Coherence Tomography
OMES: Oxic Microbial Electrosynthesis
SHE: Standard Hydrogen Electrode

## Supplementary Data

**Supplementary Figure 1:**
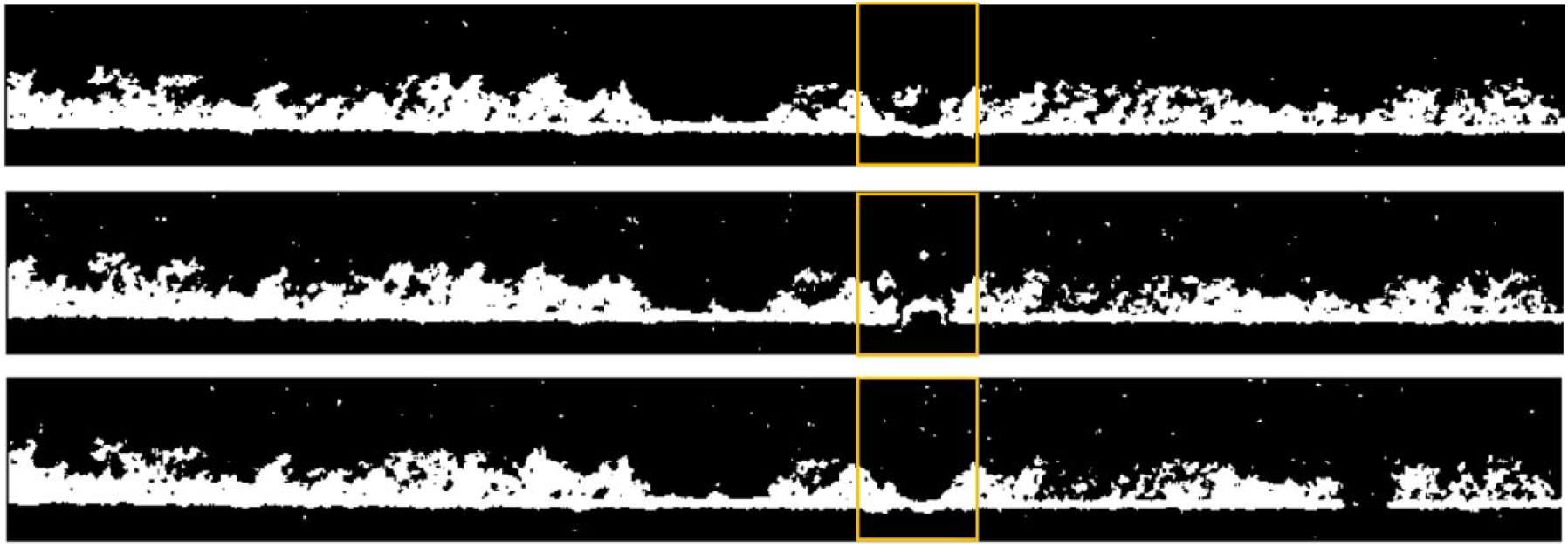
Time-resolved two dimensional OCT images of a biofilm section of 6 mm. A hydrogen bubble is evolving as can be seen in the middle image, pushing the biofilm to the side; in the bottom image the bubble is gone and detached the biofilm part above it.

## Acknowledgements

This work was supported by a grant of the Bundesministerium für Bildung und Forschung (BMBF), no. 033RC006.

## Notes

**Funding information:** This work was supported by a grant of the Bundesministerium für Bildung und Forschung (BMBF), no. 033RC006

### Competing Interest Statement

The authors have declared no competing interest.

